# A wheat kinase and immune receptor form the host-specificity barrier against the blast fungus

**DOI:** 10.1101/2022.01.27.477927

**Authors:** Sanu Arora, Andrew Steed, Rachel Goddard, Kumar Gaurav, Tom O’Hara, Adam Schoen, Nidhi Rawat, Ahmed F. Elkot, Catherine Chinoy, Martha H. Nicholson, Soichiro Asuke, Burkhard Steuernagel, Guotai Yu, Rajani Awal, Macarena Forner-Martínez, Luzie Wingen, Erin Baggs, Jonathan Clarke, Ksenia V. Krasileva, Yukio Tosa, Jonathan D. G. Jones, Vijay K. Tiwari, Brande B. H. Wulff, Paul Nicholson

## Abstract

Since emerging in Brazil in 1985, wheat blast has spread throughout South America and recently appeared in Bangladesh and Zambia. We show that two wheat resistance genes, *Rwt3* and *Rwt4*, acting as host-specificity barriers against non-Triticum blast pathotypes encode a nucleotide-binding leucine-rich repeat immune receptor and a tandem kinase, respectively. Molecular isolation of these genes allowed us to develop assays that will ensure the inclusion of these two genes in the wheat cultivars to forestall the recurrence of blast host jumps.

## Main

The occurrence of pathogen host jumps suggests that seemingly durable non-host resistance can be fragile^1^. This is illustrated by the jump of the blast fungus (*Pyricularia oryzae*, syn. *Magnaporthe oryzae*) from ryegrass to wheat in Brazil in 1985^2^. The pathogen subsequently spread to cause epidemics in other regions of Brazil and neighbouring countries including, Bolivia and Paraguay^3^. Outbreaks of wheat blast occurred in Bangladesh in 2016 and the disease was reported from Zambia in 2018^4,5^. Wheat blast is now considered to pose a threat to global wheat production^6^, and discovery and deployment of resistance genes against this pathogen are critical to mitigate its threat.

While *Pyricularia oryzae* exhibits a high level of host specificity, *Triticum* pathotypes are closely related to *Lolium* and *Avena* pathotypes^7^. Two pathogen genes, *PWT3* and *PWT4*, condition avirulence of *Avena* pathotypes on wheat (*Triticum aestivum*) while *PWT3* prevents infection of wheat by *Lolium* pathotypes^8,9^.

The resistance genes *Rwt3* and *Rwt4* in wheat recognise respectively the *PWT3* and *PWT4* avirulence gene products to prevent infection. It has been proposed that the epidemics in Brazil occurred due to the widespread cultivation of varieties lacking *Rwt3* that are susceptible to *Lolium* pathotypes^7^. *Lolium* pathotypes have also been associated with the occurrence of wheat blast in the USA^10,11^. These reports emphasize the importance of maintaining *Rwt3* and *Rwt4* in wheat cultivars to prevent future host jumps of the *Avena*, and/or *Lolium* pathotypes.

To identify candidates for *Rwt3* and *Rwt4*, we used a Triticeae bait library (Table S1, Additional File F1) to capture and sequence the NLR complements of 320 wheat lines including 300 wheat landraces from the A.E. Watkins collection harbouring the genetic diversity existing prior to intensive breeding (Table S2, Supplementary Fig. 1). We screened seedlings of the panel with Br48, a *Triticum* pathotype of *Pyricularia oryzae*, transformed with either *PWT3* or *PWT4*^7^ (Table S3; Supplementary Fig. 2, 3) and performed *k*-mer based association genetics. This led to an identification of candidate NLR genes for *PWT3* and *PWT4* recognition (Fig 1a-b, Supplementary Fig. 4, Supplementary Fig. 11) on chromosome 1D within the mapping intervals of *Rwt3* and *Rwt4*, respectively^9,12^.

**Figure 1.**
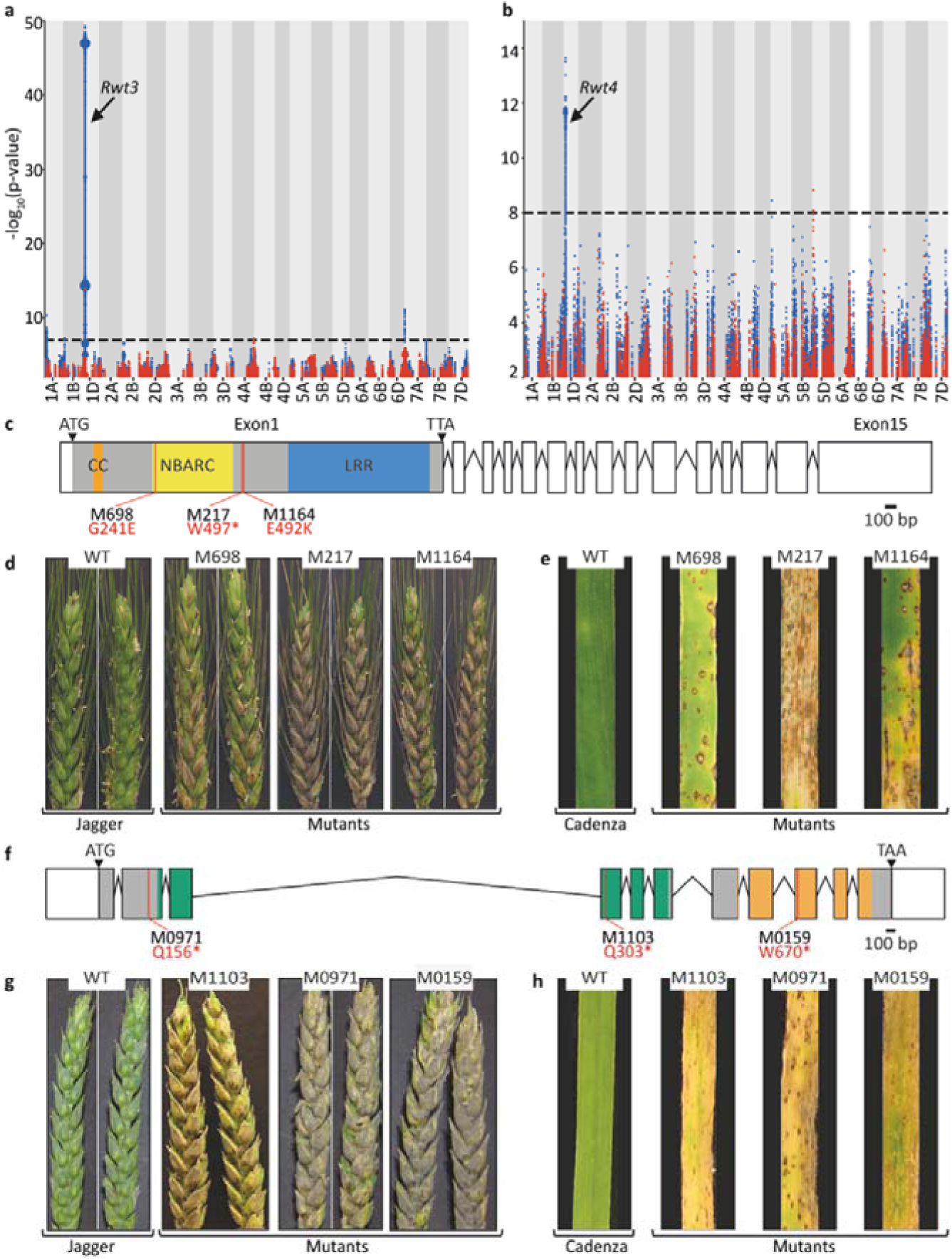
Genetic identification of the candidate genes recognising host-specific avirulence effectors of the blast fungus by *k*-mer–based association mapping on an *R*-gene enriched sequencing panel of wheat landraces. *k*-mers associated with resistance to (**a**) Br48*+Pwt3* mapped to Chinese Spring, and (**b**) Br48*+Pwt4* mapped to Jagger. Points on the y-axis depict *k*-mers positively associated with resistance in blue and negatively associated with resistance in red. Point size is proportional to the number of *k*-mers. **(c)** Structure of the NLR candidate gene for *Rwt3*. The predicted 1069 amino acids protein has domains with homology to a coiled-coil (CC), nucleotide-binding (NBARC) and leucine-rich repeats (LRR). Wheat blast **(d)** head and (**e**) detached leaf assays for the *Rwt3* Jagger mutants and wild type with Br48+*PWT3*. **(f)** Structure of the candidate gene for *Rwt4*. The predicted protein of 916 amino acids has domains with homology to a wheat tandem kinase (shown with green and orange colors). Wheat blast (**g**) head and (**h**) detached leaf assays for the *Rwt4* Cadenza mutants and wild type with Br48+*PWT4*.

Investigating the presence of these candidate genes in the NLR assemblies of *Aegilops tauschii*^13^, the D-genome progenitor of bread wheat, the *Rwt4* candidate was found only in lineage 2 (L2) while the *Rwt3* candidate was found only in lineage 1 (L1) (Table S4, Supplementary Fig. 5). This explains why we could identify only the *Rwt4* candidate, and not the *Rwt3* candidate, by phenotyping and performing association genetics on an NLR gene enrichment-sequenced *Ae. tauschii* L2 panel^13^ (Table S5, Supplementary Fig. 6, 7). The L2 origin of *Rwt4* is consistent with L2 being the major contributor of the wheat D-genome, however, the L1 origin of *Rwt3* is more remarkable considering that the L1 signature in wheat is mostly concentrated around a 5 Mb region surrounding the *Rwt3* candidate^14^ (Supplementary Fig. 8). This finding suggests that pathogen pressure could have played a significant role in post-domestication wheat evolution.

To functionally validate the *Rwt3* NLR candidate, we screened a TILLING population of Jagger^15^ and found three lines each carrying a functional mutation in this gene (Supplementary Fig. 9). One line, M217, is homozygous for a mutation causing a premature stop codon whereas another, M698, is homozygous for a mis-sense mutation (G241E) predicted to cause functional aberration in the protein (Fig. 1c, Table S6). In both the leaf and head assays of these mutants using Br48+*PWT3*, a loss of the wildtype resistance was observed (Fig. 1d-e). The third line, M1164, is heterozygous for another deleterious mis-sense mutation (E492K) (Fig. 1c, Table S6). In both the leaf and head assays of the segregating progeny of M1164 using Br48+*PWT3*, those homozygous for the mutation were found to be susceptible while the others were resistant (Fig 1d-e, Supplementary Fig. 10). The clear loss of function observed in three independently derived TILLING mutants and the co-segregation of the M1164 mutation with susceptibility shows that the *Rwt3* NLR candidate is required for resistance to *P. oryzae* expressing the *PWT3* effector.

We observed that the identified *Rwt4* NLR candidate is adjacent to an allele of a wheat tandem kinase (WTK) previously reported to confer resistance against powdery mildew^16^. The 532 kb mapping interval of powdery mildew resistance contained an allele of the *Rwt4* NLR candidate identified in our study, in addition to the WTK. On functional testing by Lu et al (2020)^16^, the WTK, and not the NLR, was found to be necessary and sufficient to confer resistance to powdery mildew and was designated as *Pm24*. Therefore, we tested both the identified *Rwt4* NLR candidate (Supplementary Fig. 11) and the linked *Pm24* allele (Supplementary Fig. 12) as candidates for *Rwt4* using the Cadenza TILLING resource^17^. For the NLR candidate, we identified four lines (two heterozygous and two homozygous) carrying mutations predicted to cause premature stop codons and six additional lines (four heterozygous and two homozygous) carrying mis-sense mutations predicted to have a significant impact on tertiary structure (Table S6). Neither the homozygous nor any progeny of the heterozygous mutants for this candidate showed an increase in susceptibility relative to the wildtype Cadenza in either leaf or head assays with Br48+*PWT4* (Supplementary Fig. 13). For the linked *Pm24* allele, we tested three lines (one homozygous and two heterozygous) carrying mutations that result in premature stop codons (Table S6). In both the leaf and head assays of the homozygous line M0159 using Br48+*PWT4*, a clear increase in susceptibility compared to the wildtype was observed (Fig. 1g-h). In the leaf and head assays of the segregating progeny of heterozygous mutants (M0971 and M1103) using Br48+*PWT4*, those homozygous for the mutation were found to be susceptible while all others were resistant (Supplementary Fig. 14). These results show that as in the case of *Pm24*, the linked WTK, and not the identified NLR candidate, is required for resistance to *P. oryzae* expressing the *PWT4* effector. The finding that *WTK* alleles, *Pm24* and *Rwt4*, are involved in resistance to two unrelated fungal pathogens suggests that it may be a broad-spectrum component of disease resistance.

We developed KASP markers for *Rwt3* and *Rwt4* (Table S7) and validated them on the core 300 Watkins accessions (Table S8). *Rwt3* is present only in 145 of the 193 Watkins accessions resistant to Br48+*PWT3* (Fig. 2a, Table S8), while *Rwt4* is present only in 136 of the 270 Watkins accessions resistant to Br48+*PWT4* (Fig. 2b, Table S8). This suggests that there are other resistance genes in the Watkins panel recognising *PWT3, PWT4* or additional effectors in Br48. We re-ran GWAS with the leaf assay disease phenotype of Br48+*PWT4*, restricted to the Watkins lines not containing *Rwt4*. Using Jagger as the reference genome, we obtained a clear peak on chromosome 1B in the region homeologous to that on 1D containing *Rwt4* (Fig. 2e), indicating that *Rwt4* has a homeologue on chromosome 1B that provides resistance to *P. oryzae* expressing the *PWT4* effector. We followed the same protocol and re-ran the GWAS with the leaf assay disease phenotype of Br48+*PWT3*, restricted to the Watkins lines not containing *Rwt3*. This identified a clear peak on chromosome 2A using Mattis as the reference genome (Fig. 2c) and another on chromosome 7A using Jagger as the reference genome (Fig. 2d). A resistance termed *Rmg2* located on chromosome 7A has previously been identified in the cultivar Thatcher^18^ and a resistance termed *Rmg7* has been reported on the distal region of the long arm of chromosome 2A of tetraploid wheat^19^. In both instances the resistances were identified using the same isolate, Br48, as used in our work suggesting that the resistances identified on chromosomes 2A and 7A may correspond to *Rmg7* and *Rmg2* reported previously. Watkins lines carrying the 7A resistance showed similar levels of resistance to both Br48 and Br48+*PWT3* (Supplementary Fig. 15) indicating that this resistance is due to interaction with Br48 and supporting its characterisation as *Rmg2*.

**Figure 2.**
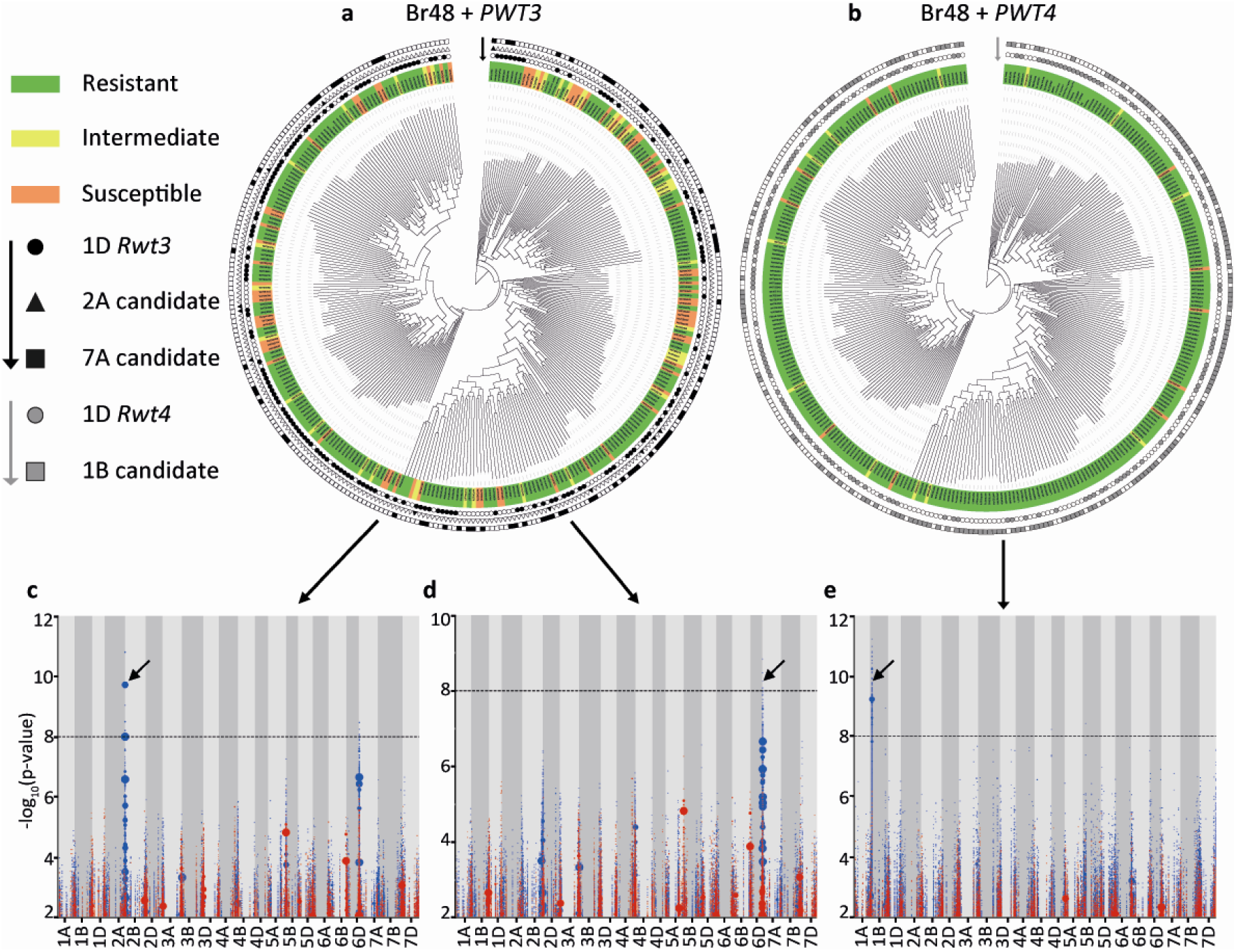
Additional resistances to blast fungus and their distribution in the diversity panel of wheat landraces. (**a**)-(**b**) *k*-mer-based phylogeny of wheat landraces showing the phenotype of an accession after inoculation with: (**a**) Br48+*Pwt3* and (**b**) Br48+*Pwt4*, and the presence of the respective candidate resistance genes. Phenotype of an accession after inoculation with a blast isolate is indicated by the color used to highlight the label of that accession, while the presence and absence of allele-specific polymorphisms is indicated by filled symbols with black/grey or white, respectively. *k*-mers significantly associated with resistance to Br48*+Pwt3* in the absence of the *Rwt3* candidate gene on chromosome 1D leads to the identification of a resistance on **(c)** chromosome 2A when mapped to the assembly of wheat cultivar SY Mattis, and (**d**) chromosome 7A when mapped to wheat cultivar Jagger. **(e)** *k*-mers significantly associated with resistance to Br48*+Pwt4* in the absence of *Rwt4* candidate gene on chromosome 1D leads to the identification of a resistance on a region of chromosome 1B containing the homeologue of *Rwt4*.

We designed a KASP-based marker for the *Rwt4* 1B homeologue (Tables S8, S9) which, along with those for *Rwt3* and *Rwt4*-1D, should enable wheat breeders to ensure that cultivars contain resistance effective against *PWT3* and *PWT4* and therefore maintain host-specificity barriers against *Lolium* and *Avena* pathotypes of *P. oryzae*. It was due to the lack of this information that *Rwt3* failed to make its way into elite cultivars such as Anahuac despite being widely present in wheat landraces (Table S9), which was the probable cause of the original wheat blast epidemic in Brazil (Fig. 3a). A future host jump of *P. oryzae* poses a high risk of host range expansion of *Triticum* pathotypes of *P. oryzae*. This risk was illustrated in the recent study of Inoue et al (2021)^20^, which showed that the resistance conferred by *Rmg8* is suppressed by *PWT4* and that the presence of *Rwt4* in wheat prevents this suppression. *Rmg8*, along with *Rmg7*, recognises the effector *AVR-Rmg8* and is one of the few reported resistances that show effectiveness against *Triticum* pathotypes of *P. oryzae* at both the seedling and head stage^21^. If *Triticum* pathotypes acquire *PWT4* from a future host jump, the resistance provided by *Rmg8* would be lost (Fig. 3b). Therefore, it is important to ensure the presence of *Rwt4* in wheat cultivars not only to prevent a future host jump but also to maintain the effectiveness of *Rmg8* against wheat blast if such an event occurs.

**Figure 3.**
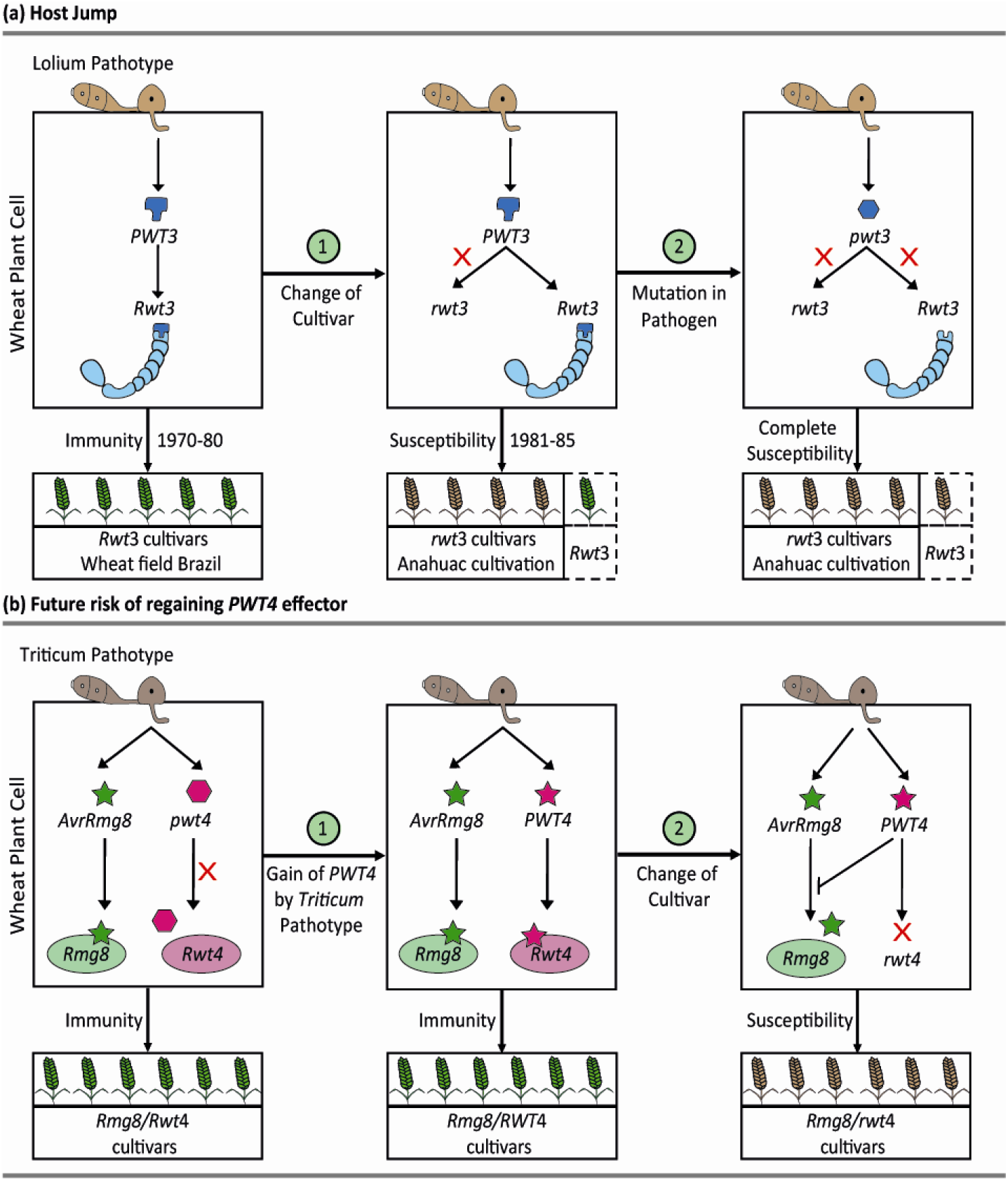
A possible working model of host jump of blast fungus from *Lolium* to wheat and a future risk associated with the reacquisition of *PWT4*. **(a)** (i) A schematically drawn wheat cell of a cultivar carrying *Rwt3* attacked by a *Lolium* isolate of the blast fungus. The *PWT3* effector is recognized by *Rwt3* thus preventing the *Lolium* isolate from infecting wheat. (ii) Widespread cultivation of cultivars lacking *Rwt3* (or having the susceptible allele, *rwt3*) allowed the *Lolium* isolate to colonize wheat. (iii) The colonizing blast population further expanded the host range by losing *PWT3* (or gaining the non-interacting effector, *pwt3*) through mutation or recombination. (**b**) (i) A schematically drawn wheat cell of a cultivar carrying *Rmg8* and *Rwt4* attacked by a *Triticum* isolate carrying *AvrRmg8*. The *AvrRmg8* effector is recognized by *Rmg8*, thus preventing *Triticum* isolate from infecting the cultivar. (ii) Gain of *Triticum* isolates gain the *PWT4* effector due to a future host jump but are still not able to infect the cultivars carrying *Rwt4* (iii) However, cultivars lacking *Rwt4* (or having the susceptible allele, *rwt4*) will be susceptible to the *Triticum* isolate carrying both *AvrRmg8* and *PWT4* even if the cultivar carries *Rmg8* because in the absence of *Rwt4, PWT4* suppresses the recognition of *AvrRmg8* by *Rmg8*. Therefore, the *Triticum* pathotype will be able to further expand its host range.

## Methods

### Watkins panel configuration

Using the SSR genotype data from Wingen et al (2014)^22^, a core set of 300 genetically diverse wheat landraces with spring growth habit were selected from the Watkins collection (Supplementary Fig. 1, Table S2) along with 20 non-Watkins lines. The DNA was extracted following a modified CTAB protocol^23^. The seeds of these lines are available from the Germplasm Resources Unit (www.seedstor.ac.uk) under Wheat Resistance gene enrichment (WREN) sequencing collection (WREN0001-WREN0320).

### Phenotyping of *Ae. tauschii* and Watkins panels with wheat blast isolates

The *M. oryzae* pathotype *Triticum* (MoT) isolate Br48 and the transformed isolates Br48+*PWT3* and Br48+*PWT4*^7^ were grown on complete medium agar (CMA). A conidial suspension of 0.3 – 0.4 x10^6^ conidia per ml was used for all inoculations. Detached seedling assays with the *Ae. tauschii* and Watkins panels were carried out as described by Goddard et al (2020)^24^ and scored for disease symptoms using a 0 – 6 scale (Supplementary Fig. 2, 3 and 6; Table S3, S5). Resistance at the heading stage was assessed according to Goddard et al (2020)^24^. Heads of *Ae. tauschii* and wheat were scored using a 0 – 6 scale (Supplementary Fig. 2e and 2f, respectively).

### Bait library design for the Watkins panel

Two bait libraries were used for the capture of the immune receptors from the Watkins panel (i) NLR Triticeae bait library V3 (https://github.com/steuernb/MutantHunter/), including 275 genes conserved in grasses^25^ and (ii) A new bait library which included NLRs extracted from the genomes of *T. turgidum* cv. Svevo and cv. Kronos and *T. dicoccoides* cv. Zavitan and only those genes that had <50% coverage by previously designed baits were used. To remove redundancies, NLR sequences were passed through CD-HIT (v4.6.8-2017-0621 -c 0.9 -G 0 - aS 0.9 -p 1). This bait design also included wheat domestication genes *VRN1A* (AY747598), *Wx1* (AY050174), *Q* (AY702956), *Rht-b1* (JX993615), *Rht-d1* (HE585643), *NAM-B1* (MG587710) as well as wheat orthologs of known immune signalling components ICS1, NPR1, NDR1, EDS1, PAD4, SRFR1, SAG101, RAR1, SGT1, HSP90.2, HSP90.4, RIN4, ADR1 and PBS1 extracted through BioMart (Table S1, Additional File 1). The bait probes were designed by Arbor Bioscience and filtered with their Repeat Mask pipeline which removed the baits that were >50% Repeat Masked and any non-NLR baits with >3 hits in the wheat genome. To balance for the low copy number genes, baits derived from domestication genes were multiplied 10x and those derived from immune signalling genes were 3x compared to the baits derived from NLRs.

### Library construction and sequencing of the Watkins panel

Illumina libraries with an average insert size of 700□bp were enriched by Arbor Biosciences, Michigan, USA, as previously described^26^, and sequenced on an Illumina HiSeq with either 150 or 250 PE reads at Novogene, China to generate an average of 3.82 Gb per accession (Table S2). The raw reads were trimmed using Trimmomatic v0.2^27^ and *de novo* assembled with the CLC Assembly Cell (http://www.clcbio.com/products/clc-assembly-cell/) using word size (-w□=□64) with standard parameters.

### Generating Watkins *k*-mer presence/absence matrix and its phylogeny

A presence/absence matrix of *k*-mers (k□=□51) was constructed from trimmed raw data using Jellyfish^28^ as described in Arora et al (2019)^13^. *k*-mers occurring in less than four accessions or in all but three or fewer accessions were removed during the construction of the matrix. From the *k*-mer matrix generated with Watkins RenSeq data, 5310 randomly extracted *k*-mers were used to build a UPGMA (unweighted pair group method with arithmetic mean) tree with 100 bootstraps.

### *k*-mer based association mapping

For the reference genomes of *T. aestivum* - Chinese Spring^29^, Jagger and Mattis^30^ – and of *Ae. tauschii* AY61^31^, NLRs were predicted using NLR-Annotator^32^ and their sequences along with 3kb sequence from both upstream and downstream region (if available) were extracted using samtools (version 1.9) to create the corresponding reference NLR assemblies. The disease phenotypes were averaged across the replicates after removing the non-numerical values and the mean phenotype scores multiplied by −1 so that a higher value represents a higher resistance. For those *k*-mers of a reference NLR assembly whose presence/absence in the panel correlates with the phenotype, that is, the absolute value of Pearson’s correlation obtained was higher than 0.1, a p-value was assigned using linear regression while taking the three most significant PCA dimensions as covariates to control for the population structure. A stringent cut-off of 8, based on Bonferroni-adjustment^14^ to a *p*-value of 0.05, was chosen for Watkins RenSeq association mapping, while a cut-off of 7 was chosen for *Ae. tauschii* L2 RenSeq association mapping (Supplementary Fig. 7).

### *In silico* gene structure prediction

The *Rwt3* NLR candidate gene transcript is 5,937 bp. Only one of the 15 annotated exons (grey colored exon in Fig. 1c) appears to be translated into protein. This exon encodes a protein of 1069 amino acids with a coiled-coil domain, an NB-ARC domain and several leucine rich repeats (LRRs) motifs at the C-terminus (Supplementary Fig. 4). The *Rwt4* NLR candidate gene is 3,117 bp with three exons. The predicted protein of 1038 amino acids contain domains with homology to a coiled-coil (CC) domain, two NB-ARC domains and two LRR at the C-terminus (Supplementary Fig. 11). The *Rwt4* WTK candidate has an open reading frame of 2,751 bp which has eleven predicted exons that encode a protein of 916 amino acids with putative tandem protein kinase domains (Fig. 1f; Supplementary Fig. 12). Domains were predicted by NCBI and Pfam databases. The gene structure of both *Rwt3* and *Rwt4* NLR candidate genes was consistent with that predicted using cDNA RenSeq data of Watkins lines.

### Identification and phenotyping of Cadenza TILLING mutants to test the function of *Rwt4*

Cadenza TILLING lines^17^ for the NLR candidate for *Rwt4* were identified within the Plant Ensembl database for the gene TraesCS1D02G059000 (http://plants.ensembl.org/Triticum_aestivum/Gene/). Lines containing mutations leading to premature stop codons and those for which the ‘sorting intolerant from tolerant’ (SIFT) score was 0.0 or 0.01 were selected for phenotyping. For the *Rwt4* kinase candidate gene, Cadenza TILLING lines were identified for the gene TraesCS1D02G058900. Details of the mutations present in the Cadenza TILLING lines is provided in the Table S6.

### Identification and phenotyping of Jagger TILLING mutants to test the function of *Rwt3*

For selecting mutations in the *Rwt3* candidate gene (TraesCS1D02G029900), TILLING was performed in wheat cultivar Jagger^15^ using genome specific primer pairs (Supplementary Fig. 9a-d). The effects of the mutations on the predicted protein were analysed using SnapGene® software (version 5.0.7 from GSL Biotech). The effects of missense mutations were determined using PROVEAN (Protein Variation Effect Analyzer) v1.1 software^33^. Selected lines were phenotyped as described above. Details of the mutations is provided in Table S6.

### KASP analysis and sequencing of TILLING lines to confirm mutations

Kompetitive Allele-Specific PCR (KASP) (LGC Genomics) was performed to confirm mutations where suitable PCR primers could be designed. Alternatively, the region containing the mutation was amplified and purified products were sequenced by Eurofins Genomics. Sequence analysis was performed with Geneious Prime software.

### Anahuac DNA preparation, sequencing, and assembly to check presence of *Rwt3* gene

To confirm that Anahuac is a non-carrier of *Rwt3*, we captured its NLR complement using the bait libraries described above. The *Rwt3* NLR candidate was absent in the CLC assembly generated as described above.

### KASP marker design to detect *Rwt3* and *Rwt4* in wheat cultivars and Watkins collection

The regions differentiating resistant and susceptible alleles of *Rwt4* from the *Ae. tauschii* L2 panel were used to design KASP markers. The KASP marker discriminated between resistant and susceptible accessions in *Ae. tauschii* L2 panel but did not distinguish reliably between resistant and susceptible lines in the Watkins panel. The resistant allele of *Rwt4* was the same in both the *Ae. tauschii* L2 and the Watkins panels but the susceptible allele of *Rwt4* in the Watkins panel originated from *Ae. tauschii* L3 and not *Ae. tauschii* L2. This is consistent with the multi-lineage hybridisation hypothesis proposed in Gaurav et al 2021^14^. A new marker was designed by comparing the common resistant allele with susceptible alleles from both the *Ae. tauschii* L2 and Watkins panels (Table S7) that successfully distinguished between the resistant and susceptible alleles in the wheat lines (Table S8).

We used the D-genomes of 11 chromosome-scale wheat assemblies^30^ to fetch the D-genome susceptible allele of *Rwt3* and designed KASP markers (Table S7). The marker distinguished resistant from susceptible lines and had a high correlation with presence-absence scored with *in silico* markers (Table S8). KASP markers were tested on the entire Watkins panel (∼900) to understand the distribution of these genes in the landrace collection (Table S9).

### Characterisation of the resistance identified on chromosome 7A

A set of Watkins lines were genotyped as carrying either *Rwt3* or the 7A resistance or having neither or both resistances. All accessions were phenotyped in leaf assays using isolates Br48 and Br48+*PWT3*. Accessions lacking either resistance were susceptible to both isolates (Supplementary Fig. 15). Accessions carrying either the 7A resistance alone or both the 7A resistance and *Rwt3* showed similar level of resistance to both Br48 and Br48+*PWT3*.

## Acknowledgements

The high-performance computing resources and services used in this work were supported by the Norwich Bioscience Institutes Partnership (NBIP) Computing infrastructure for Science (CiS) group alongside the Earlham Institute (EI) scientific computing group. We are grateful to the John Innes Centre (JIC) Horticultural Services for plant husbandry; EI for providing open access to the Kronos genome. This research was financed by the Biotechnology and Biological Sciences Research Council (BBSRC) Designing Future Wheat Cross-Institute Strategic Programme to BBHW and PN (BBS/E/J/000PR9780); a John Innes Centre Institute Strategic Grant to BW; Science, Technology & Innovation Funding Authority (STDF), Egypt-UK Newton-Mosharafa Institutional Links award, Project ID (30718) to AFE and BBHW; the Gordon and Betty Moore Foundation through grant GBMF4725 to the Two Blades Foundation; and the Gatsby Charitable Foundation to JDGJ; National Science Foundation (Award#1943155) and USDA NIFA (Award#2020-67013-32558 and 2020-67013-31460) to NR and VT; European Research Commission grant (ERC-2016-STG-716233-MIREDI) to KVK and BBSRC Norwich Research Park Doctoral Training Grant (BB/M011216/1) for supporting EB.

## Author contributions

This work was conceived by PN, JC and BBHW. Watkins panel configuration, DNA extraction and sequence acquisition (BBHW, LW, MFM, RA, SA, GY, AFE, JDGJ). Bait library design (KVK, EB, BS), *k*-mer matrix construction and association mapping (SA, KG), candidate genes discovery and analysis (SA, KG), Phylogenetic analysis (SA, KG), Blast isolates (YT, SAs), Phenotyping of diversity panels and TILLING mutants (AS, RG, TH, PN, CC, MHN), KASP marker design and analysis (SA, AS, KG, PN), Jagger mutants identification (VT, ASc, NR), Mutant confirmation and segregation (AS, RG, PN), cDNA RenSeq data (SA, AFE), Drafted manuscript (SA, PN, KG, RG, AS, VT, YT, ASc, NR, KK) and designed figures (SA, PN, KG, RG, AS, ASc, LW).

