## Supplementary Figures for "A wheat kinase and immune receptor form the host-specificity barrier against the blast fungus"

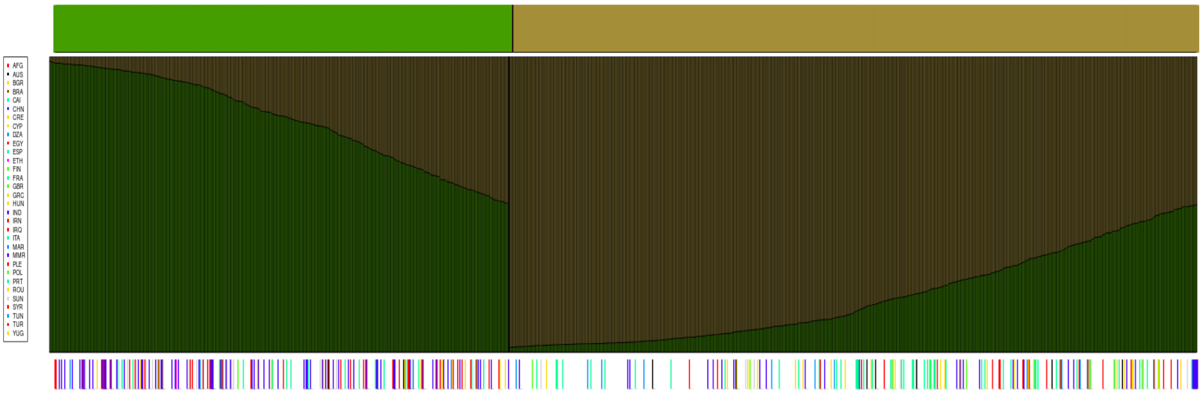


**S1** STRUCTURE assignment of the landraces in the Watkins collection (1054 landraces) from Wingen et al (2014). The top row shows the assignment to a top-level split into two large ancestral populations. The middle row indicates the proportion of ancestral characteristics of each landrace cultivar (LCs). The bottom row colour code of country/region of origin is only shown for the 314 members of the core set. Unselected LCs are shown in white. Colour codes group countries into geographic regions.

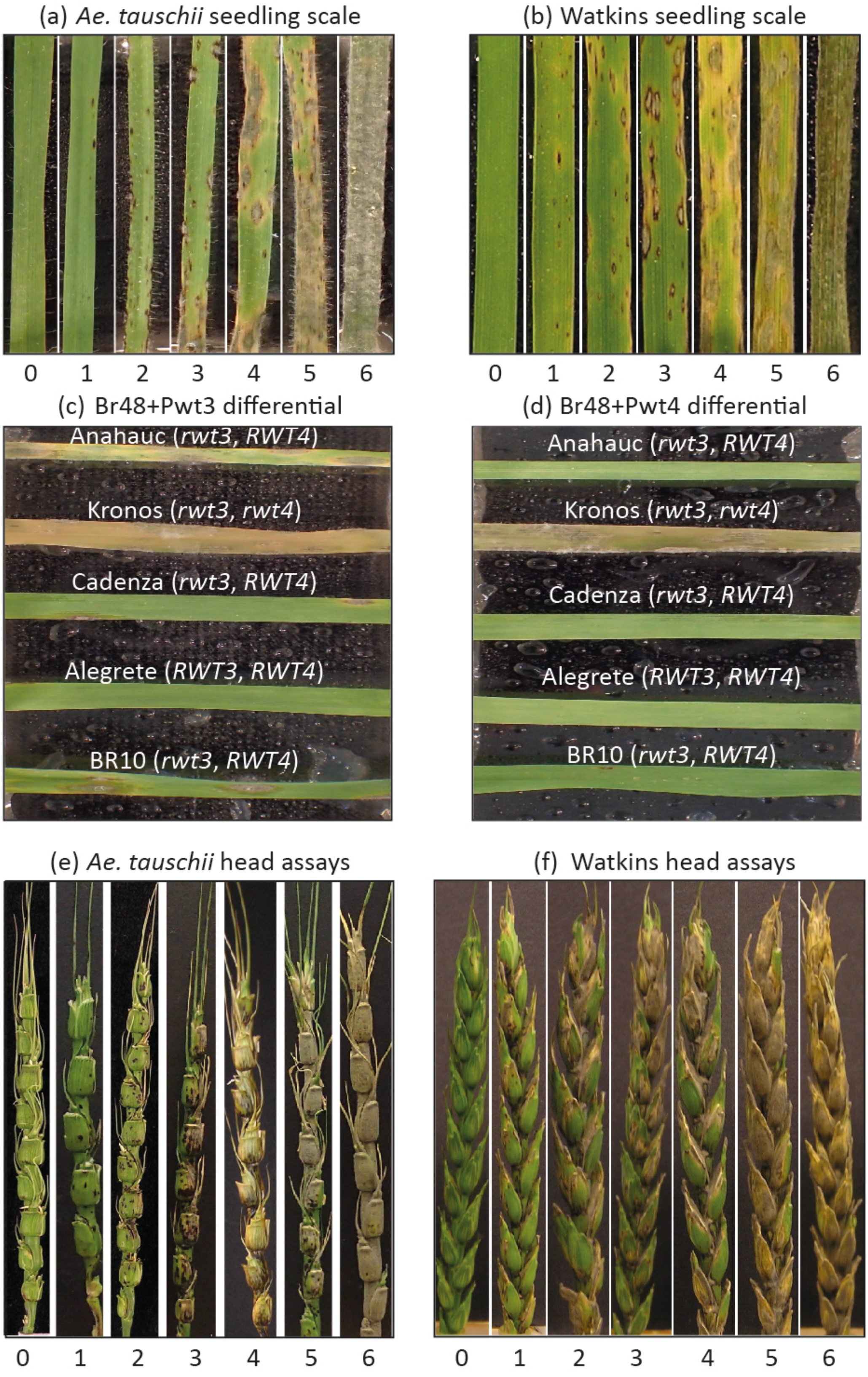


**S2** Wheat blast detached leaves scoring scale (0-6) at 6dpi for (a) *Ae. tauschii* and (b) Watkins panel. Variation observed for the differential lines (Anahuac, Kronos, Cadenza, Alegrete, BR10) upon phenotyping with (c) Br48+*PWT3* and (d) Br48+*PWT4*. Phenotype scale (0-6) at head stage for (e) *Ae. tauschii* and (f) Watkins panels.

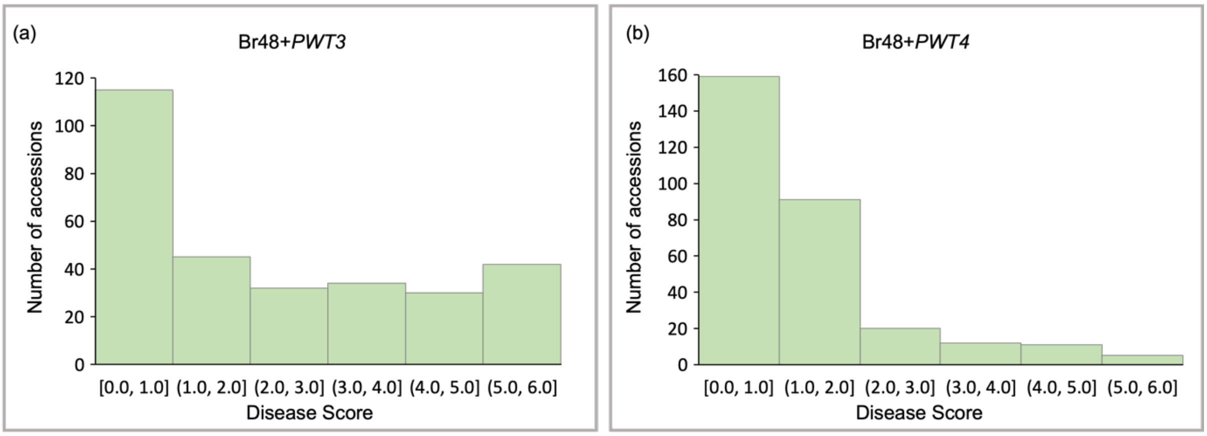


**S3** Bar graphs showing phenotypic variation observed in the Watkins panel for (a) Br48+*PWT3* and (b) Br48+*PWT4*.

(a)

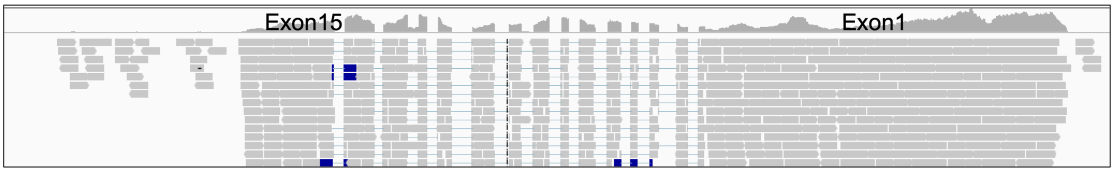


(b)

```
MAAALGAAQLLLGKVLTKLSDERMASYVASSELGLDSQKIKDDIMYTVGLLQQAQGRGTSDIPGLTDLLVKLSQKADEAEK
VLNELRYFMIRDQINGTQLVKPDLGACLKTKKGHARHTIGNCLPFIFRPCTSSQQDGTMSDGHDHVDELPFDSVSISIKIK
SVLEETHSICVHVSDLLKLIPNHGSSSTTATTVTRIRPTRGSMVAQDTMYGRRDIFEKTIHAITSCGEILSVLPVVGPGGI
GKTTLIQHLYNDKRIEEHFDVRIWVCVSIDFAVLKLTREIFRCIIATEEEGRNSAPSETASIEQLQESIAQRLQSKRFLIV
LDDVWQLNSEDEWHTLLAPFKKGGAKGNMVLVTTRFPSIAGRVKTVDPVELQGLDTDDFFAFFEECIFGGLDKPMHYRDEL
TDIARDIVDKLKGSPLAAKTVGRLLRKDLSWEHWTRVLESMEWENKQSDNDTMPALYLSYDYLPFLLQKCFSYLSLFPEDH
RFSHSEINRFWIAVGIIDSNHPGNNNCLEELVDRGFLRRVHGLFGEEYYVIHDLLHELSRSVSTQECLNISVLDFKADTVP
PSIQHLSITLENKYDGIFLEEISKLKIKIDIANLRTLMIFSAYEERIAGILKDTFEEVDSLHVLFIVVKSLDDLPKGFSKL
IHLQYLKLGSPIGIEMALPSTLARFYHLKFLDLKDWHGSSNVPKDISHLVNLQDFIAKKELHSSVLEVGKMKYLRELKQFC
VKKESVGFELRELGELTELGGELRICNLENVATKEEASEAKLMSKRNLKKLTLVWGRKQSTIDGDVLDALQPHPDLRELRI
ANHGGAVGPSWLCVDMLLKQLGVLHLEGLSWDTLPPFGQLPRLTKLILMRISGVHQLALLSNLTSLTHLTLQDCVKLTVDG
FNPLTTVNLKVLVVFNCRWDRSCPESVAADLLTKVASSRVMPAGSFRLEQLKVDSISAVLVTPICNLVAHTLQTLIFCHDH
RIKSFTEEQEKALLLLTSLRHLTFDGCGALQSLPRGLNRLSSLKDLEVLWCPEMGSIPKEGFPVSLEILRIRPCSPEVREQ
IEKLRRTSPGLSVRYE
```

**S4** (a) Mapping of RNA-Seq reads to the *Rwt3* NLR candidate gene present in the Chinese Spring genome and (b) the predicted amino acid sequence of *Rwt3* gene with coiled-coil (orange), NB-ARC (yellow) and LRR (blue) domains.

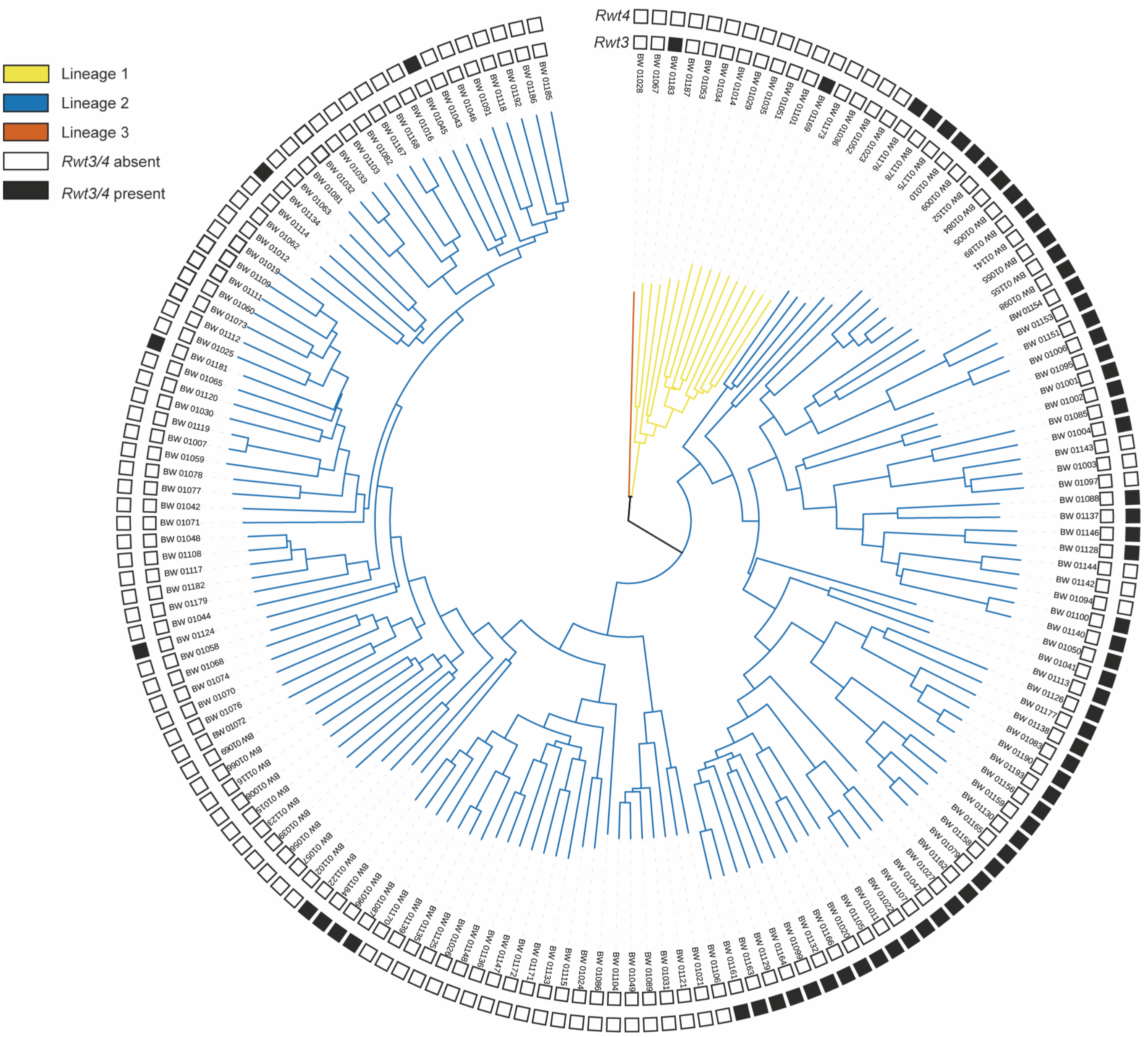


**S5** Presence of *Rwt3* and *Rwt4* candidate resistance genes in the phylogeny of the *Ae. tauschii* panel (adapted from Arora et al 2019).

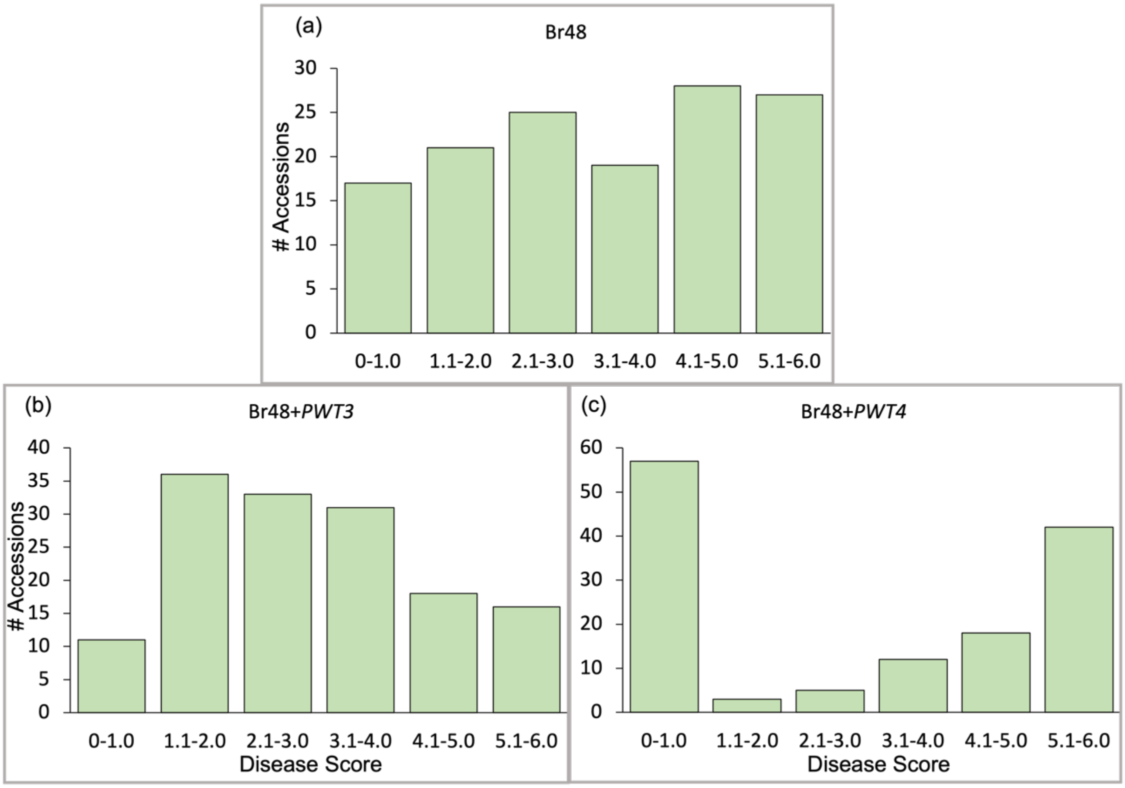


**S6** Bar graphs showing phenotypic variation observed in the *Ae. tauschii* panel for (a) Br48 (b) Br48+*PWT3* and (c) Br48+*PWT4*.

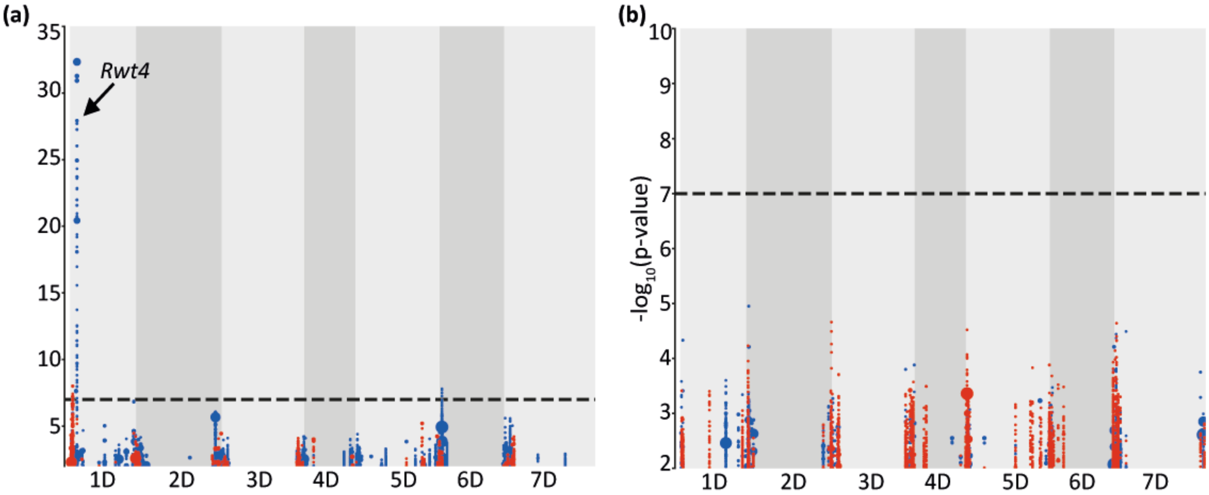


**S7** Association mapping plots showing (a) identification of a clear peak in the *Ae. tauschii* panel screened with Br48+*PWT4*, however, (b) no clear association was observed for *PWT3* recognition in this panel.

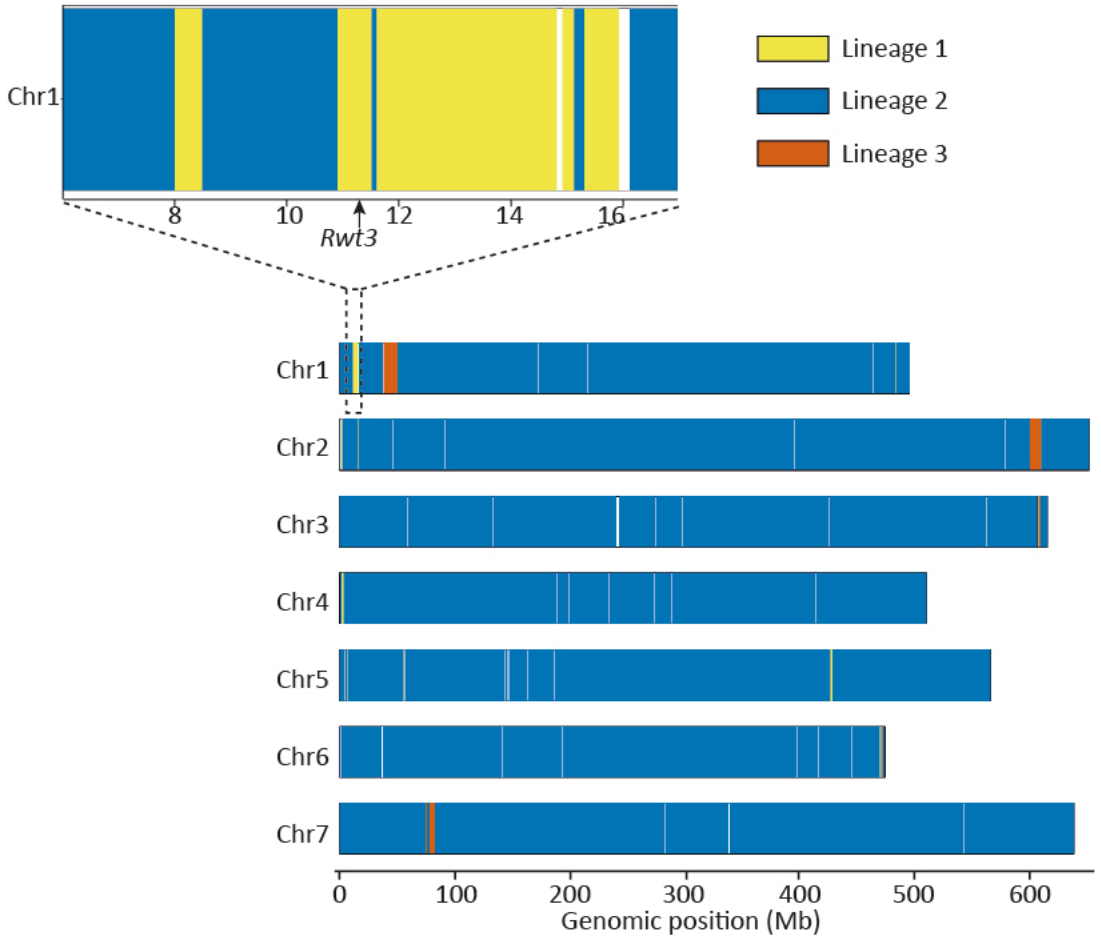


**S8** The pattern of lineage-specific contribution to the Chinese Spring D-subgenome across all the seven chromosomes (adapted from Gaurav et al 2021). The genomic region most enriched with lineage 1 contribution is located on chromosome 1DS surrounding *Rwt3*.

(a)

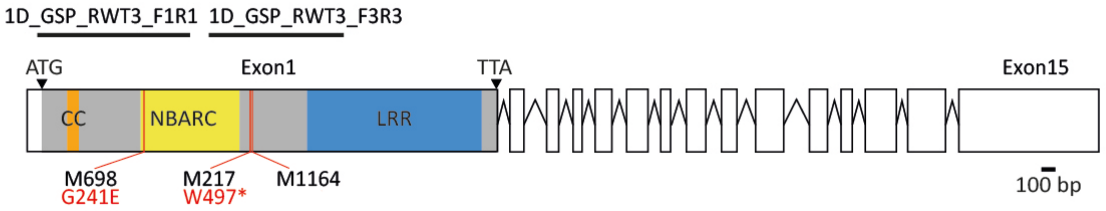


(b)

| Primer name | Sequence | Product length (bp) |
|---|---|---|
| 1D_GSP_RWT3_F1 | CTACTGTCCATCGCACTACCG | 990 bp |
| 1D_GSP_RWT3_R1 | CTGTTTCACTGGGAGCACTATTT | |
| 1D_GSP_RWT3_F1 | TTCTGGATGATGTGTGGCAACT | 773 bp |
| 1D_GSP_RWT3_R1 | GGTGGGACTGTGTCAGCTT | |

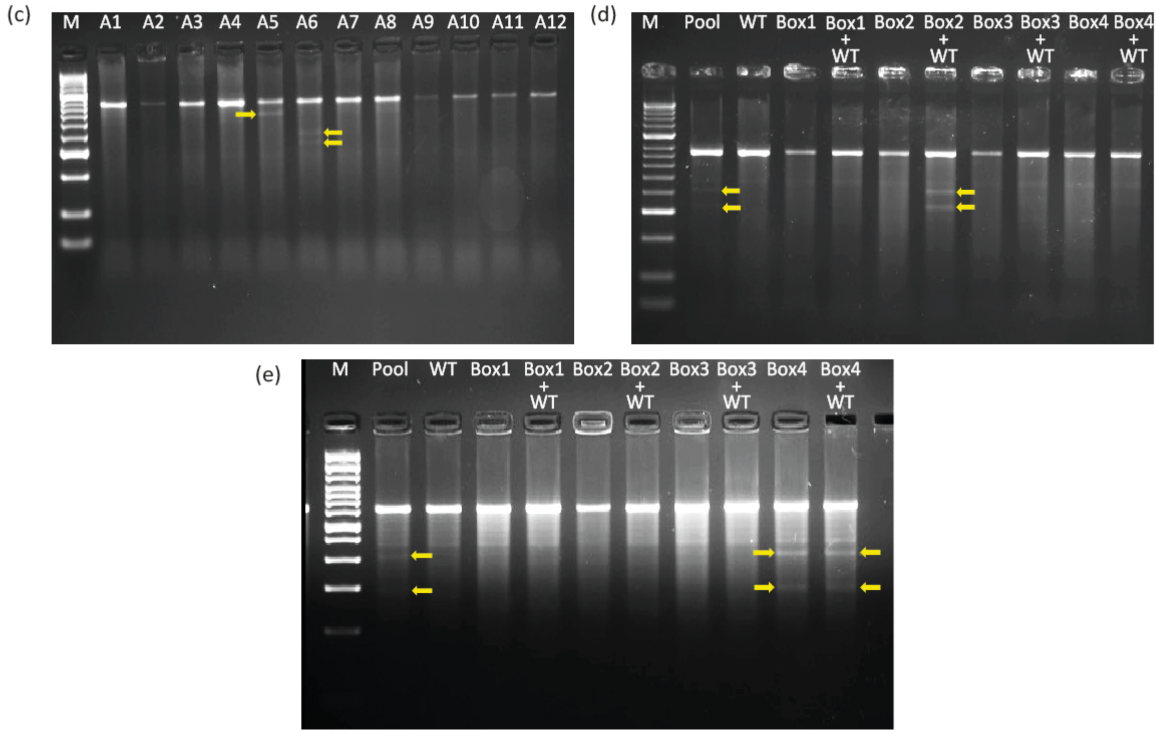


**S9** *Rwt3* Jagger mutant selection (a) Position of the two primer pairs in the *Rwt3* gene selected for performing TILLING and location of the mutants on the gene selected for disease phenotyping. (b) Primers used for TILLING the Jagger mutagenized population (c) Identification of mutants in a row of 12 pools of the Jagger TILLING population, Deconvolution and zygosity determination of: (d) a homozygous mutant and (e) a heterozygous mutant.

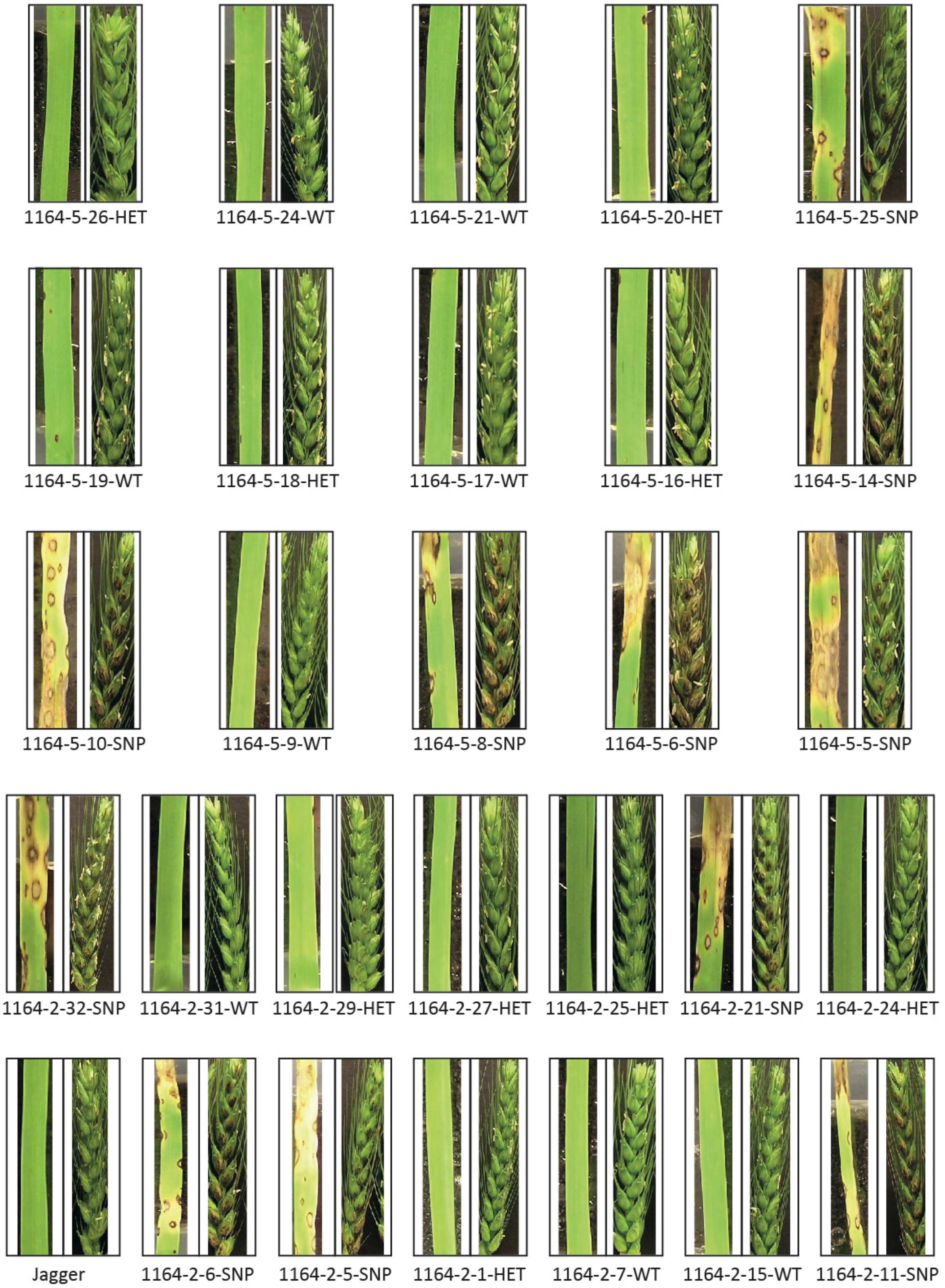


**S10** Leaf and head assays of the segregating progeny of Jagger for *Rwt3* heterozygous mutant 1164 using Br48 + *PWT3*.

(a)

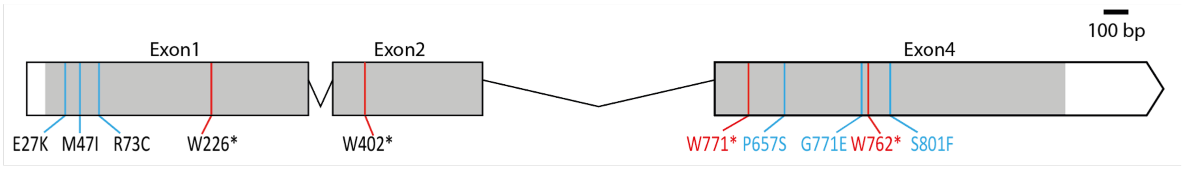


(b)
MEEIVLGLSKTVVEGTLVKVKAAIDEEAKLKLAVQSDLVFITGEFEMMQSFLNVADAERIKNNAVKTWVRQLRDLAYDTEDCIELVVHLDP
KPRWWRRLLVLPCLPAVSLPMDNAAAEIKELKDRVEFVSQRNMRYKLITDFGGAKSVAQQQLDRAAGALDIAVEEAAKKGRSLV**DLTELIP**
**RMEDRPELGVISVWGTGGDLGVASIVRKAYDDPEICKNFHSRGWAKLTHPFDPRKILRSLFIQFCTNTAAPQKQGETLDVDSLLRMEKTVV**
**EEGELVKQFVSHVSTHRFLVVLEGVSTMAEWDALRMYLPDMGNGSQIIVSTQHFDIASLCTGQLHKVSELRKFTPDHSVCVFFKEV**SGGDD
PGNAMLEGDDET**ELIGRDTEEDELMEQIDEVSRVISVWGIAGVGKSALVRSAYCKYLQDRPNFAGLNLTYIHGRVNVPHPFNLREFCRSLL**
**LNLSSEPAKTKEDAVAELANMTNPIDVCRKLLSDDQQQKQCLIVIDDLQSTEEWDRIKQALSFGKECCIIVVTNDSSVAKYCSGQDQKLMF**
**NVKGLGDEHALDLFNTVYDGVGNHITRDEDMINQSKLMLNKCGGLPKVIVAIGCFLAKSLNWEAMNTNFIHQLENNPELASVHSMFSW**LDS
YFHNCPDELKPCIFYLSIFPRRHGIRRRRLVRRWIAEGYSRDTDGNLAEVNGENYFSRLVNLSMLTDAERDVNVAGTATGAGTGRRMAMCN
VNDFFREYIVSRRMEENHVFALEGRCSQTTRRTGRHLVIDKSWDGDESVFNRIEFSRLRSVTVFGPWKPFLASDKMRVLRVLDLEGTEGLT
DNDIKNIVKRLPRLKFLSLRGCKHIFRLPKSLGRLRQLQTLDVRHTAVAGLPATVVKLQKLQYVRAGNTPEEYAVSDSTESSTSVPTCFSC
GPVGNHIVGVEVPRGVENLTNLHTLGVIKATVAGLKELKKLTQLRKLGVSGINRRNHKELCAVISGHGHLESLSIWFDKDTERIADPCSPS
NDDFKPPEKLRRLKLHGHTDNLPAWVSTLKIFVTQDQ*

**S11** The intron-exon structure of the NLR candidate gene for *Rwt4.* The predicted 1038 amino acids protein has domains with homology to a coiled-coil (orange), nucleotide-binding (yellow) and leucine-rich repeats (blue).

MGGYEFQRAELDALEGVVRDPTAEPMSLTLPLLRHITNDFSPEFEISKDDSAVVYLGVLPSGFRVAVKKL
HNRHWLEDEDAFINEVSIAMKAAKKNTVRVIGYCQHTQEQIAEYEGKQVFAELTERLICTEYVPNGPLSG
HIEGKICAQMDGYEFQRAELDALERVVRDTSAEPMSLTLPLLRHITNDFSDESRIGRGGFAVVYLGVLPS
GLRIAVKRLSNIAYMNESAFQNEVFITMKATHKNTVRFMGYCSQIQGKLIEHNGQHVFAQLEERLICVEY
APKGTLDAHIGDYGELDWNQRYQILKGICQGLHHLHDEMHVFHGDIKPANILIGDNLVPKIYDFGLSQMF
EEEETERIVKNMAGTLCYLAPEVLNTHMMSFKAEIYSLGVVIEELLTGKKGWIDEDVRKQLKGLRKTLVK
EGAFSSWENKYHQVRTCMEIGQDCIDPNPHKRPTLFEIIQRLNEAEDMNYSAASLWQVQSGDEESDLSDT
EALETETTSEFLPSDEEPASVGKTGETSTQEPDKPDLVSKLPASVDLSDLKVLEKITDDFSHERIVGKDG
TFKGCHKAFVYKGDIPRREMIAVKRLIGVEIPVEKFKREAEQFIRLDHKNIVKVASYCHDQSTGHRLVQF
KGKPLPQLFQGPEQLLCYEYMHNGSLRDYLMGQGSREIDWQMRYKLIKGTCAGLDYLHKGRAGCPIVHLN
LSPSNVLLDHNYIPRITGFDFSKLIGEKNTKSVVLKLNGPIAYLPPDFFHSKGTDLKYLATVDIYSLGLM
ILEIATQQEIKGIHGVLIKSIEENWREESQITRLYTSLGADELRQVKMCIDIGLDCVKSNPEKRPTAGAI
MLWLDKESKPVPVSRAGAGVLPRPPVPTNINHAGRIQGIPIEKSQVLIPSFACNEYSSVKIQNLHLLLFM
FIQVDQ

**S12** Protein sequence of *Rwt4* WTK gene with two (green and orange coloured) predicted kinase domains.

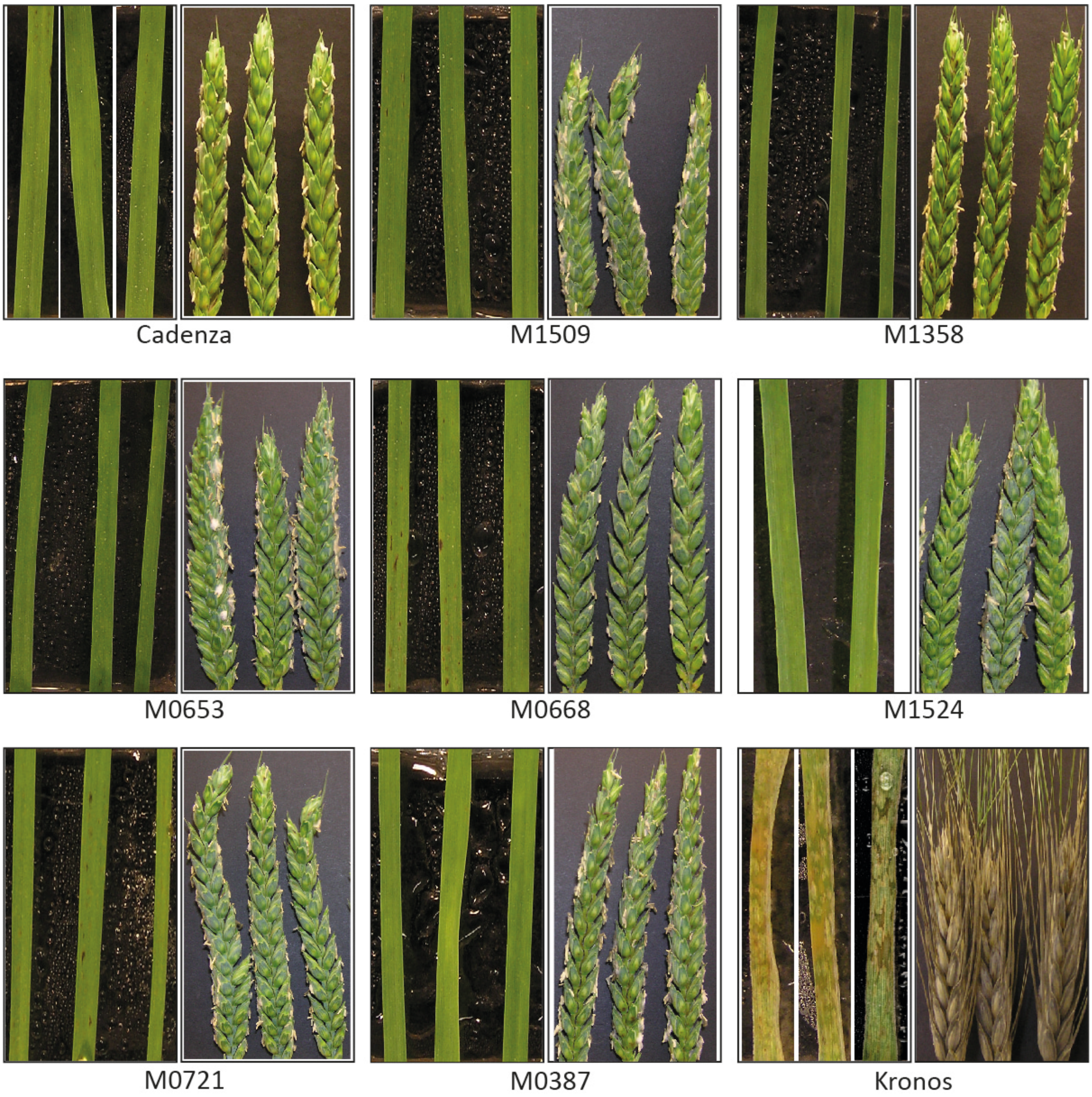


**S13** Phenotyping of the mutants for the *Rwt4* NLR candidate gene with Br48+*Pwt4* at both seedling and head stage.

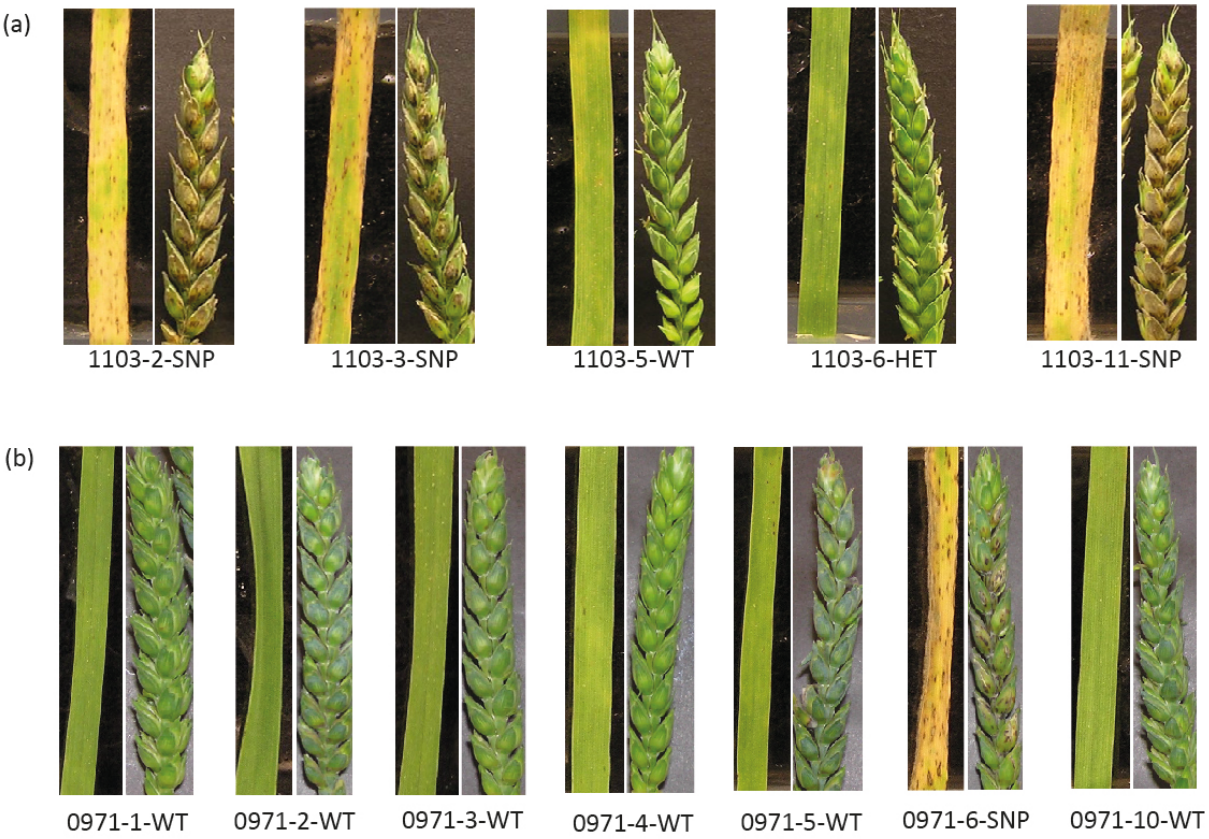


**S14** Genotype and phenotype correlation of the segregating progenies of *Rwt4* WTK mutants - Cadenza1103 and Cadenza 0971.

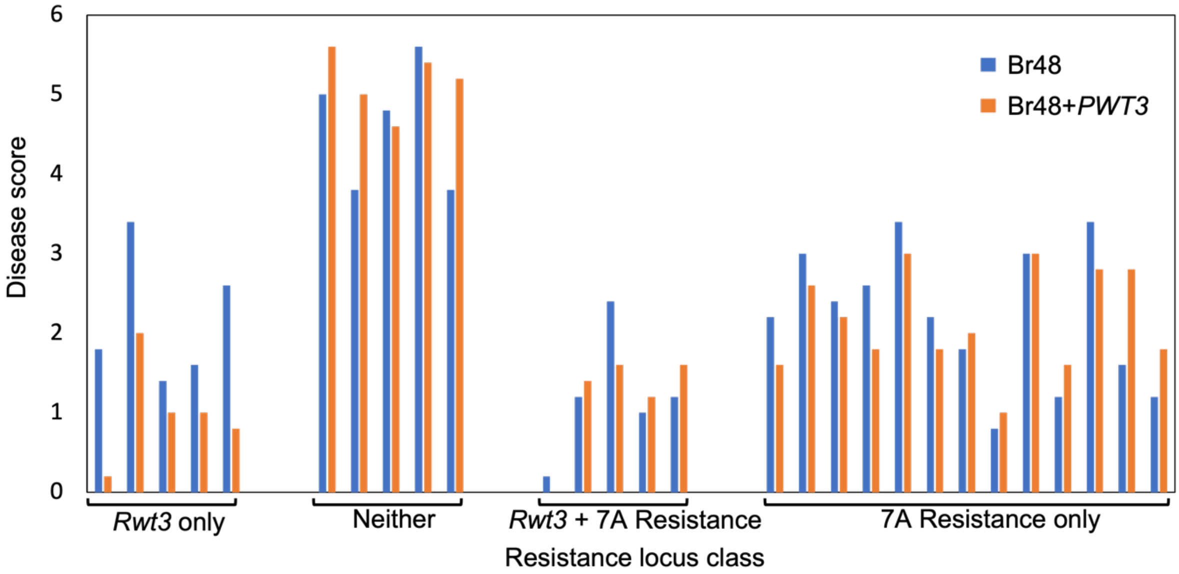


**S15** Effect of the presence of *Rwt3* and the resistance locus identified on chromosome 7A on the resistance to Br48 and Br48+*PWT3*.

### Supplementary Tables (refer to the attached excel sheet for the tables)

**S1** Gene IDs of additional sequences used to design new baits.

**S2** Metadata for the 300 core Watkins lines and amount of RenSeq data received for each line

**S3** Phenotype data for the Watkins panel screened with Br48+*PWT3* and Br48+*PWT4*

**S4** *In silico* distribution of *Rwt4* and *Rwt3* NLR candidates identified on wheat chromosome 1D in the *Ae. tauschii* panel

**S5** Phenotype data for the *Ae. tauschii* panel screened with Br48, Br48+*PWT4* and Br48+*PWT3*

**S6** Summary of the *Rwt3* and *Rwt4* (NLR and WTK candidates) TILLING mutants including their phenotype and mutation information.

**S7** KASP diagnostic markers designed for *Rwt4-1D, Rwt4-1B* and *Rwt3*

**S8** The distribution of *Rwt4*-1D and 1B, *Rwt3,* and the resistance loci identified on chromosomes 2A and 7A in the Watkins panel inferred using *in silico* markers. Additionally, the panel was genotyped with KASP markers for *Rwt3* and *Rwt4*-1D and a high correlation was observed between KASP and *in silico* markers

**S9** Distribution of *Rwt3*, *Rwt4*-1B and *Rwt4*-1D markers in 943 accessions of the Watkins collection.

**Additional File**
**F1** Sequence of NLRs extracted from the genomes of *T. turgidum* cv. Svevo and cv. Kronos and *T. dicoccoides* cv. Zavitan for use in the bait library.
